# DenVar: Density-based Variation analysis of multiplex imaging data

**DOI:** 10.1101/2021.09.27.462056

**Authors:** Souvik Seal, Thao Vu, Tusharkanti Ghosh, Julia Wrobel, Debashis Ghosh

## Abstract

Multiplex immunohistochemistry (mIHC) and multiplexed ion beam imaging (MIBI) platforms have become increasingly popular for studying complex single-cell biology in the tumor microenvironment (TME) of cancer subjects. Studying the intensity of the proteins that regulate important cell-functions, often known as functional markers, in the TME becomes extremely crucial for subject-specific assessment of risks, such as risk of recurrence and risk of death. The conventional approach requires selection of two thresholds, one to define the cells of the TME as positive or negative for a particular functional marker, and the other to classify the subjects based on the proportion of the positive cells. The selection of the thresholds has a large impact on the results and an arbitrary selection can lead to an incomprehensible conclusion. In light of this problem, we present a threshold-free distance between the subjects based on the probability densities of the functional markers. The distance can be used to classify the subjects into meaningful groups or can be used in a linear mixed model setup for testing association with clinical outcomes. The method gets rid of the subjectivity bias of the thresholding-based approach, enabling an easier but interpretable analysis of these types of data. With the proposed method, we analyze a lung cancer dataset from an mIHC platform, finding the difference in the density of functional marker HLA-DR to be significantly associated with the overall survival. The approach is also applied on an MIBI triple-negative breast cancer dataset to analyze effects of multiple functional markers. Finally, we demonstrate the reliability of our method through extensive simulation studies.

## 1 Introduction

In recent years, various technologies are being used for probing single-cell spatial biology, for example, multiparameter immunofluorescence (Bataille *and others*, 2006), imaging mass cytometry (IMC) (Giesen *and others*, 2014; Chang *and others*, 2017; Ali *and others*, 2020), multiplex immunohistochemistry (mIHC) (Halse *and others*, 2018; Tan *and others*, 2020; Vu *and others*, 2021) and multiplexed ion beam imaging (MIBI) (Angelo *and others*, 2014; Seal *and others*, 2021b). These technologies, often referred to as multiplex tissue imaging, offer the potential for researchers to explore the bases of many different biological mechanisms. Multiplex tissue imaging platforms such as Vectra 3.0 (Akoya Biosciences) (Huang *and others*, 2013), Vectra Polaris (Akoya Biosciences) (Pollan *and others*, 2020), MIBI (Ion-path Inc.) (Keren *and others*, 2019; Ptacek *and others*, 2020) produce images with similar structure. In particular, each image is two dimensional, collected at cell- and nucleus-level resolution and proteins in the sample have been labeled with antibodies called “markers” that attach to cell membranes. Typically, mIHC images have 6-8 markers, whereas MIBI images can have more than 40 markers.

Majority of the above markers are surface or phenotypic markers, such as CD4, CD3, CD8, CD68 etc. (Jondal *and others*, 1972; Zola *and others*, 2007; Shipkova and Wieland, 2012) which are primarily used for cell type identification. Additionally, there are several functional markers including HLA-DR (Jendro *and others*, 1991; Oczenski *and others*, 2003; Saraiva *and others*, 2018), PD-1, PD-L1, Lag3 etc. (Nguyen and Ohashi, 2015; Han *and others*, 2020; Phillips *and others*, 2021) that dictate or regulate important cell-functions. Both surface and functional markers are quantified as continuous valued marker intensities. For a phenotypic marker, a threshold is drawn to indicate whether a cell is positive or negative for the particular marker. Then one or more of these binarized phenotypic markers are used to classify the cells into different types based on biological knowledge of marker co-expression. With the functional markers, the interest lies in finding out if abundance or over-expression of the markers across the cells of the tumor microenvironment (TME) (Whiteside, 2008; Binnewies *and others*, 2018) have significant impact on subject-level clinical outcomes, such as survival or recurrence (Sahlberg *and others*, 2014; Koguchi *and others*, 2015; Johnson *and others*, 2020). A two-step thresholding-based approach (Bulian *and others*, 2014; Costa *and others*, 2017; Missassi *and others*, 2021) is typically used in this context which we describe next.

The two steps in the thresholding-based approach involve identifying cells positive for a marker and classifying patients into different groups according to the proportions of positive cells. The group labels can be used in a linear regression framework to test association with the outcomes of interest (Chen *and others*, 2016; Chang *and others*, 2018; Yang *and others*, 2019). For example, Johnson *and others* (2021) defines the cells to be positive for HLA-DR (also known as, MHCII) if the corresponding mean marker intensity is greater than 0.05. Next, they classify the subjects into two groups, MHCII: High and MHCII: Low if the proportion of cancer cells positive for HLA-DR is greater or smaller than 5% respectively. Finally, they test if these two groups of subjects have different 5-year overall survival. Instead of grouping the subjects based on the proportion of positive cells, another approach would be to directly test if the vector of the proportion of positive cells is associated with the outcome (Patwa *and others*, 2021).

The aforementioned thresholding-based method clearly requires judicious selection of the cut-offs that greatly influence the subsequent steps of the analysis (Harris *and others*, 2021). The result is bound to vary for different thresholding values; and a poor choice of thresholds may produce an uninformative and uninterpretable result. There is a plethora of helpful guidelines on choosing these thresholds in different contexts (Barnett *and others*, 1999; Kimball *and others*, 2018; Cossarizza *and others*, 2019; BIO-RAD, 2021). However, there is no universal solution or rule of thumb. Thus, the method remains prone to subjectivity bias and lacks robustness.

In this paper, we propose a threshold-free method for distinguishing the difference between the subjects with respect to the functional markers. We treat the expression of every marker as a continuous random variable having realizations in the cells of a subject. For every marker, we compare its probability distribution or equivalently, density between every pair of subjects. Our exact algorithm is as follows. First, for every subject, the probability density of each marker is estimated using kernel density estimation (KDE) (Silverman, 1981; Sheather and Jones, 1991; Ghosh *and others*, 2006). Next, a density based distance (Basu *and others*, 1998; Jones *and others*, 2001) known as Jensen-Shannon distance (Endres and Schindelin, 2003; Fuglede and Topsoe, 2004) is used to quantify the difference in the estimated density across the subjects. The matrix of distances between subjects for every marker can then be used to classify them into meaningful groups using hierarchical clustering (Murtagh, 1985; Murtagh and Legendre, 2014). In a linear regression framework, the cluster-labels can be tested for association with clinical outcomes. The distance matrix can also be directly used in a linear mixed model (Hoffman, 2013; Seal *and others*, 2021a) or equivalently, a kernel machine regression framework (Liu *and others*, 2008; Hua and Ghosh, 2015; Ge *and others*, 2016; Jensen *and others*, 2019). Using our proposed method, we have analyzed an mIHC dataset on lung cancer (Johnson *and others*, 2021) from the University of Colorado School of Medicine, finding out that the difference in HLA-DR marker density in tumor cells is associated with 5-year overall survival of the subjects. We have also applied the proposed method on a publicly available triple negative breast cancer (TNBC) dataset (Keren *and others*, 2018) from the MIBI platform finding the density of an immunoregulatory protein, PD-1 to have significant effect on overall survival. We have performed extensive simulation studies mimicking the characteristics of the real datasets to check the reliability and robustness of our method.

## 2 Materials and Methods

Suppose there are *M* functional markers and *N* subjects with *j*-th subject having *n_j_* cells. Let *X_kij_* denote the scaled expression, between 0 and 1, of *k*-th marker in *i*-th cell of *j*-th subject for *k* = 1,2,…, *M*, *i* = 1,2,…, *n_j_* and *j* = 1, 2,…, *N*. Let *Y* (*N* × 1 vector) be a subject-level outcome of interest and *C* be an *N* × *p* matrix of *p* subject-level covariates.

### 2.1 Traditional thresholding based approach for clustering subjects

To study if abundance of marker *k* is associated with a subject’s survival or any other outcome of interest (*Y*), the conventional approach is to classify the subjects into two or more groups using a thresholding based approach. First, consider a threshold *t*_1_ and check how many of the *n_j_* cells of subject *j* have marker expression more than that threshold i.e. the number of cells with *X_kij_* > *t*_1_. Such cells are referred to as the cells positive for marker *k*. The proportion of the cells positive for a marker *k* in subject *j* is denoted as, 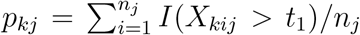, where *I*(.) is the indicator function. Another threshold *t*_2_ is chosen to classify the subjects into two groups, one with subjects more than *t*_2_% positive cells i.e. subjects with *p_kj_* > *t*_2_, and the other with subjects less than *t*_2_% positive cells i.e. subjects with *p_kj_* < *t*_2_. Then, test if these two groups of people have differential rate of survival (or, associated with some other outcome of interest). This can easily be extended to allow more than two groups.

Denote the clustering variable as *Z_kj_* ≡ *I*(*p_kj_* > *t*_2_) with *Z_kj_* being a binary variable taking values zero and one. When *Y* is a continuous/categorical outcome, a standard multiple linear regression model with **Z**_*k*_ = (*Z*_*k*1_,…, *Z_kN_*)^*T*^ as a predictor can be written as

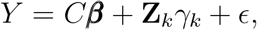

where ***β***, *γ* are fixed effects and *ε* is an *N* × 1 error vector following multivariate normal distribution (MVN) with mean **0** and identity covariance matrix 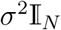. After estimating the parameters, the null hypothesis, *H*_0_: *γ_k_* = 0, can be tested using the Wald test (Gourieroux *and others*, 1982).

Next, we consider the case of *Y* being a survival or recurrence outcome. Let the outcome of the *j*-th individual be *Y_j_* = *min*(*T_j_*, *U_j_*), where *T_j_* is the time to event and *U_j_* is the censoring time. Let *δ_j_* ≡ *I*(*T_j_* ≤ *U_j_*) be the corresponding censoring indicator. Assuming that *T_j_* and *U_j_* are conditionally independent given the covariates for *j* = 1, 2,…, *N*, the hazard function for the Cox proportional hazards (PH) model (Andersen and Gill, 1982; Lin and Wei, 1989; Therneau and Grambsch, 2000) with fixed effects can be written as,

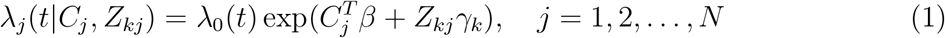

where λ_*j*_(*t*|*C_j_*, *Z_kj_*) is the hazard of the *j*-th subject at time *t*, given the vector of covariates *C_j_* and the cluster label *Z_kj_* and λ_0_(*t*) is an unspecified baseline hazard at time *t*. To test the null hypothesis: *H*_0_: *γ_k_* = 0, a likelihood ratio test (LRT) (Therneau, 1997) can be considered. The above procedure can be conducted individually for *k* = 1,…, *M* and the influential markers can be reported.

As pointed out earlier, the biggest difficulty with this approach lies in choosing the thresholds, *t*_1_ and *t*_2_ appropriately. In most cases, one would run the approach for different pairs of (*t*_1_, *t*_2_) and choose the one that leads to the most interpretable result. Thus, the step of threshold-selection remains entirely subjective and the results are bound to vary largely depending on the selected thresholds.

### 2.2 Proposed Method: Distance based clustering using marker probability density of subjects

To avoid the bias inherent in the thresholding-based approach, we propose a distance between the subjects based on each marker k that would be devoid of subjectivity and can easily be tested for association with a outcome of interest. First, we discuss the concept of divergence or distance between two probability distributions and then, implement it in the context of our problem.

#### 2.2.1 Jensen-Shannon distance

Let 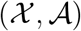 be a measurable space (Billingsley, 2008) where 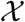 denotes the sample space and 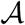 the *σ*-algebra of measurable events. Consider a dominating measure *μ* and denote the set of probability distributions as, 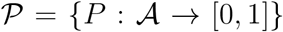. In this context, the Jensen-Shannon distance (JSD) (Endres and Schindelin, 2003; Fuglede and Topsoe, 2004; Nielsen, 2019) between two probability distributions, 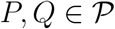 can be defined as,

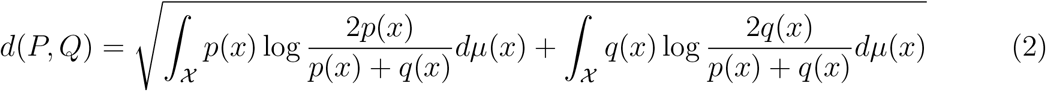

where *p*, *q* are the Radon-Nikodym derivatives or densities (Nikodym, 1930) of *P* and *Q* with respect to a dominating measure *μ*. Unlike other divergences between distributions, such as Kullback-Leibler divergence (Van Erven and Harremos, 2014), the Jensen-Shannon distance (JSD) satisfies the properties of being a metric (Lawvere, 1973) between probability measures. To formalize this, a metric 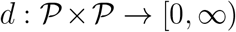 satisfies the following three axioms:

1. Identity: *d*(*P*, *Q*) = 0 iff *P* = *Q*,
2. Symmetry: *d*(*P*, *Q*) = *d*(*Q*, *P*),
3. Triangle Inequality: *d*(*P*, *Q*) + *d*(*Q*, *R*) ≥ *d*(*P*, *R*) where 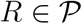.

Note that, *P* = *Q* implies *p*(*x*) = *q*(*x*) almost everywhere w.r.t *μ* (Athreya and Lahiri, 2006; Feller, 2008). JSD can be shown to be bounded above by 2log(2) and bounded below by 0 (Endres and Schindelin, 2003). JSD has been used in many different areas, such as bioinformatics (Sims *and others*, 2009), social sciences (DeDeo *and others*, 2013), and more recently, in generative adversarial networks (GANs) (Goodfellow *and others*, 2014), a popular technique in deep learning.

#### 2.2.2 Formulation of the distance in our context

For every subject *j*, we assume that the expression of marker *k* is a continuous random variable, denoted by *X_kj_*, taking values between 0 and 1. *X_kj_* is observed in *n_j_* cells as, *X*_*k*1*j*_, *X*_*k*2*j*_,…, *X_kn_j_j_*. Let the probability distribution function and the density function of *X_kj_* be denoted by, *F_kj_* and *f_kj_* respectively. Next, we consider the set-up described in Section 2.2.1 with 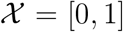 and 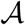 being the corresponding *σ*-algebra of measurable events. Then the set, 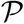 contains the distribution functions, *F_kj_* for *j* = 1,2,…, *N* and *k* = 1,2,…, *M*. Finally, using Equation 2, the distance between two subjects (*j*, *j*′) in terms of the probability distribution of marker k can be quantified as,

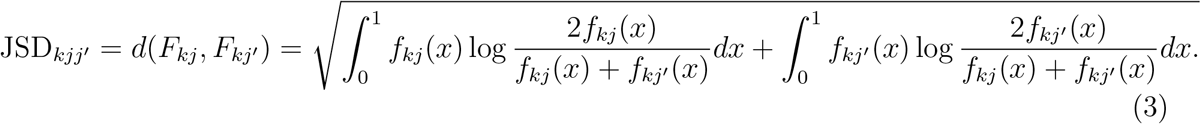

A large value of JSD_*kjj*′_ will imply that there is a clear difference in the distribution or equivalently, density of *k*-th marker between the pair of subjects, (*j*, *j*′). A small value will imply that the distributions are close. The distance matrix between all the subjects based on *k*-th marker can then be constructed as, JSD_*k*_ = [[JSD_*kjj*′]_].

In real data, the density function *f_kj_* will be unknown. Therefore, we compute corresponding kernel density estimate (KDE) 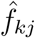 (Silverman, 1981; Sheather and Jones, 1991; Ghosh *and others*, 2006) using the observations: *X_kij_*’s for *i* = 1,…, *n_j_*. 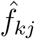 typically has the form: 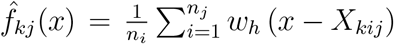, where *w_h_* is a Gaussian kernel with bandwidth parameter *h*, chosen using Silverman’s rule of thumb (Silverman, 2018). Using the KDEs, JSD_*kjj*′_ from Equation (3) can be estimated as,

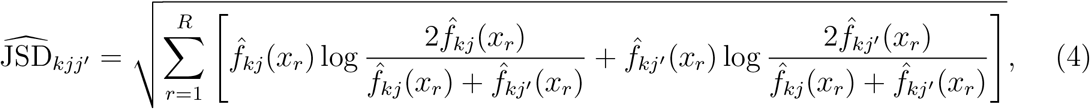

where *x_r_*, *r* = 1,…, *R* are grid-points in the interval [0,1]. In our simulations and real data analysis, we keep *R* at 1024 and choose equidistant grid-points. We have noticed that for *R* ≥ 512, the results do not alter. We make sure that the estimated densities integrate to 1 by appropriately scaling them.

#### 2.2.3 Using the distance in association analysis

Next, we construct suitable tests for testing the association of the distance matrix with dependent variable, *Y*.

##### Test based on hierarchical clustering

The estimated distance matrix 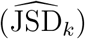 can be subjected to hierarchical clustering (Murtagh, 1985; Murtagh and Legendre, 2014) for classifying the subjects into two or more groups. Suppose, we obtain a vector of cluster labels: **Z**_*k*_ = (*Z*_*k*1_,…, *Z_kN_*)^*T*^. Then, exactly the same models, described in Section 2.1 and corresponding tests, can be used to determine if the differential expression of the *k*-th marker is associated with *Y*.

##### Test based on linear mixed model

The distance matrix can be transformed into a similarity matrix (Vert *and others*, 2004) as, 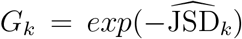. When *Y* is a continuous/categorical outcome, *G_k_* can be incorporated in a linear mixed model framework, particularly popular in the context of heritability estimation (Hoffman, 2013; Seal *and others*, 2021a), as,

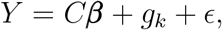

where ***β*** is the vector of fixed effects, *g_k_* = (*g*_*k*1_, *g*_*k*2_,…, *g_kn_*)^*T*^ is the vector of random effects following 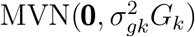 and *ε* is an error vector following 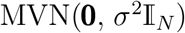. The null hypothesis: 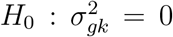 can be tested using a likelihood ratio test (Crainiceanu and Ruppert, 2004). Note that, such a linear mixed model setup has been shown to be equivalent to a kernel machine regression framework by Liu *and others* (2008). In a standard kernel machine regression framework, there is one additional width parameter, *ρ* that has to be estimated.

Next, we consider the case of *Y* being a survival or recurrence outcome. Using the same definitions and conditional independence assumptions of *T_j_*, *U_j_* and covariates as in Section 2.1, the hazard function for the Cox proportional hazards (PH) model with random effects (Therneau *and others*, 2015) can be written as,

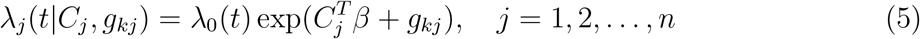

where λ_*j*_(*t*|*C_j_*, *g_kj_*) is the hazard of the *j*-th subject at time *t*, given the vector of covariates *C_j_* and the random effect *g_kj_* and λ_0_(*t*) is an unspecified baseline hazard at time *t*. To test the null hypothesis, 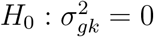, an LRT based on integrated partial likelihoods (Therneau and Therneau, 2015) can be considered. However, it is to be kept in mind that usually a large sample size is needed to obtain a precise estimate of the random effect variance (Maas and Hox, 2005; Bell *and others*, 2010; Austin and Leckie, 2018). The problem would possibly be exacerbated in the Cox PH model with random effects because the partial likelihood would depend on the number of events (Peduzzi *and others*, 1996; Vittinghoff and McCulloch, 2007; Kocak and Onar-Thomas, 2012; Ogundimu *and others*, 2016). Therefore, we do not recommend using this test unless the sample size is sufficiently large.

## 3 Results

We first discuss the application of our method on the real datasets. We analyzed two datasets: an mIHC lung cancer dataset (Johnson *and others*, 2021) and an MIBI breast cancer dataset (Keren *and others*, 2018). The first dataset has a single functional marker, HLA-DR and the second dataset has four immunoregulatory proteins, PD-1, PD-L1, Lag3 and IDO. We applied the method proposed in Section 2.2 on both the datasets. In all the analyses, the markers were scaled to have expression value between 0 and 1.

### 3.1 Application to mIHC Lung Cancer data

In the mIHC lung cancer dataset, there are 153 subjects each with 3-5 images (in total, 761 images). The subjects have varying number of cells identified (from 3,755 to 16,949). The cells come from two different tissue regions: tumor and stroma and are classified into either of the six different cell types: CD14+, CD19+, CD4+, CD8+, CK+ and Other, based on the expression of phenotypic markers, CD19, CD3, CK, CD8 and CD14. A functional marker, HLA-DR (also known as MHCII), is also measured in each of the cells. Using the thresholding-based approach described in Section 2.1, Johnson *and others* (2021) classified the subjects into two groups, a) MHCII: High and b) MHCII: Low based on the proportion of CK+ tumor cells that are also positive for HLA-DR. They found out that there is significant difference in 5-year overall survival between the groups. Analogously, we were interested in answering the question: whether 5-year overall survival of a subject is associated with the HLA-DR density in CK+ tumor cells. We first computed the JSD matrix between the subjects as discussed in Section 2.2.2 based on the density of HLADR marker in CK+ tumor cells. Next, we performed a hierarchical clustering using the computed JSD matrix to classify the subjects into two groups. Next, we tested if there is a difference in survival between the subjects of the two groups using the test based on the Cox PH model with fixed effects described in Equation 1. Figure 1 shows the Kaplan-Meier curves (Efron, 1988) of the two groups of subjects. We noticed that Hazard Ratio (HR) is large (> 2) and the *p*-value is significant (< 0.015) indicating that 5-year overall survival is associated with the probability density of HLA-DR in CK+ tumor cells. Figure 2 shows individual and mean HLA-DR probability density of different subjects from the two clusters. We noticed that the individual densities from cluster 1 were more right-skewed compared to those from cluster 2 which led to the mean density of cluster 1 having very high mode compared to that of cluster 2. We also checked the degree of conformity between Johnson *and others* (2021)’s classification and the classification based on our method. Table 1 displays the comparison between the classifications. Accompanying values of Rand index (RI) and adjusted Rand index (ARI) were respectively, 0.64 and 0.29 which made us conclude that the classifications moderately agreed with each other. Figure 3 shows individual and mean HLA-DR probability density of the subjects from groups, MHCII: High and MHCII: Low. We noticed that some of the subjects from MHCII: High group actually had density functions similar to the average density of MHCII: Low group meaning that the thresholding-based method was incapable of fully capturing the differences between the density profiles.

**Figure 1:**
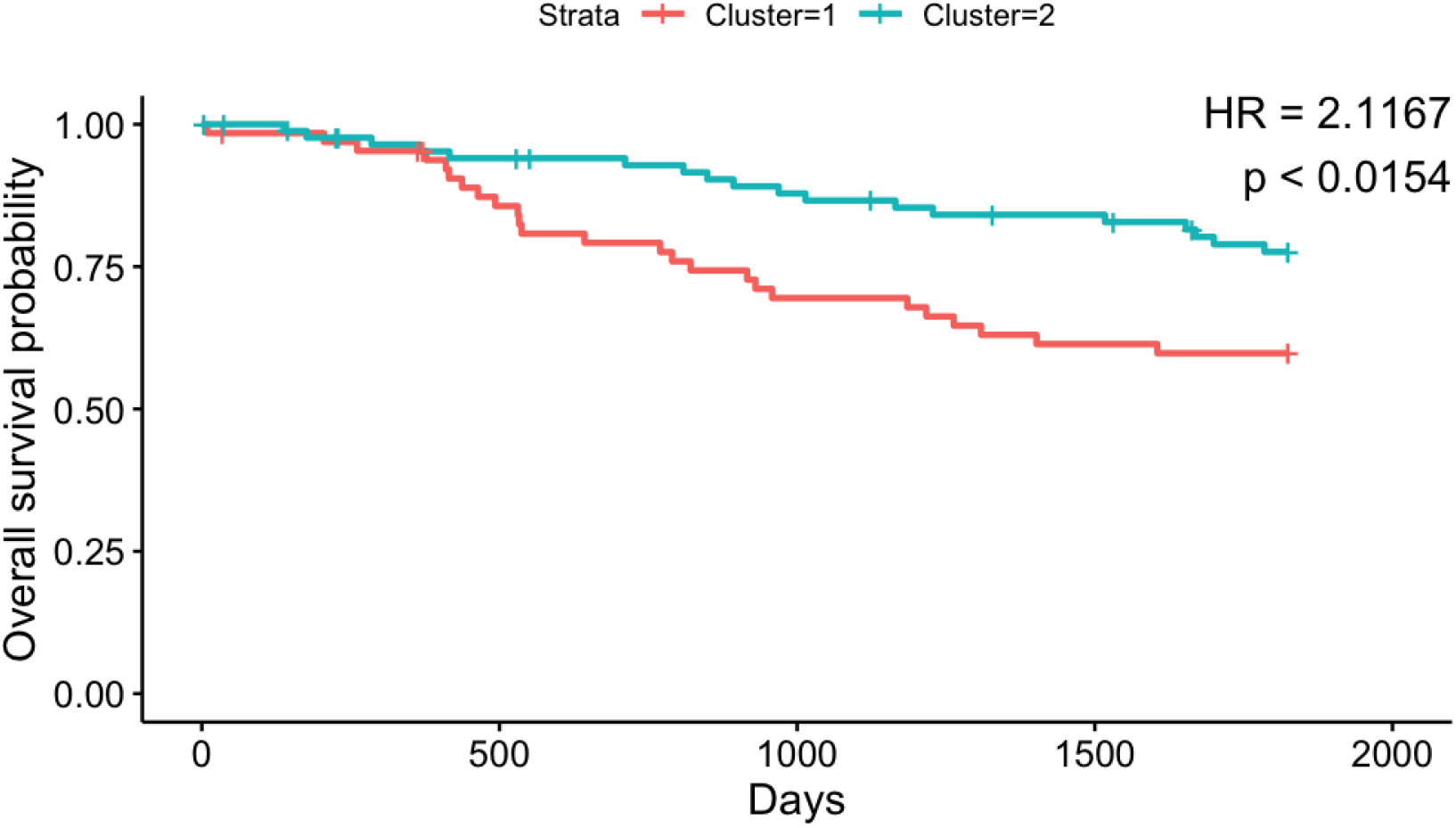
KM curves of 5-year overall Survival of 153 subjects from the lung cancer dataset, color coded by the clusters found comparing HLA-DR marker density in CK+ Tumor cells. Also, displayed are the Hazard ratio (HR) and the p-value corresponding to the test, *H*_0_: *γ* = 0 from Equation 1. Notice that HR is large (> 2) and the p-value is significant as well indicating that the two clusters have significant difference in survival probability.

**Figure 2:**
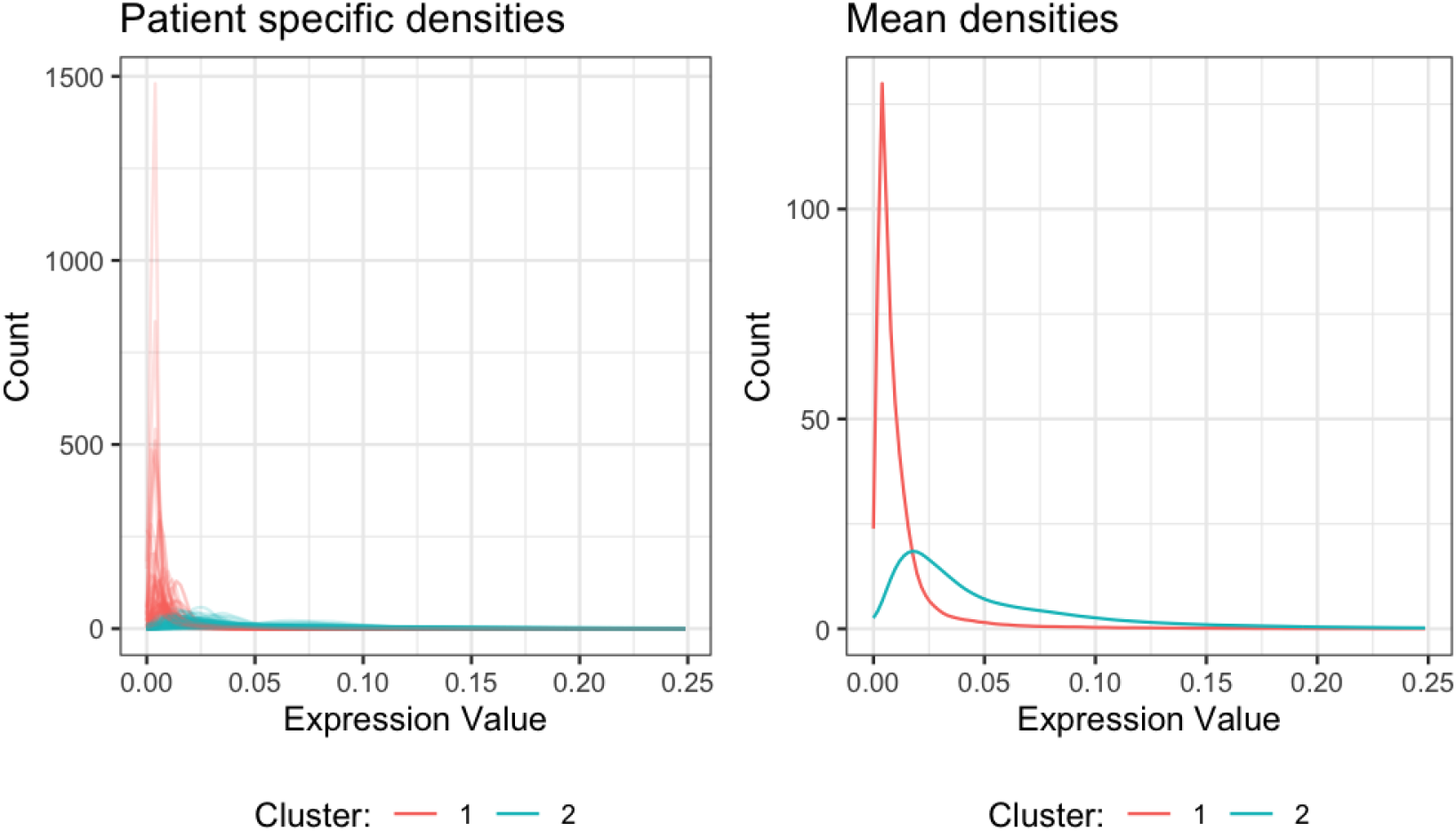
Individual (on the left) and mean (on the right) HLA-DR marker probability density (in CK+ tumor cells) of the subjects from the two clusters found using JSD based clustering proposed in Section 2.2. Notice that the individual densities from cluster 1 are more right-skewed than the densities from cluster 2. Consequently, the mean density of cluster 1 is also more right-skewed than that of cluster 2 and has a much higher peak.

**Table 1:** Number of subjects common between the groups found using the thresholding-based method and our proposed method in the lung cancer dataset.

|  | Cluster 1 | Cluster 2 |
| --- | --- | --- |
| MHCII: High | 80 | 17 |
| MHCII: Low | 18 | 38 |

**Figure 3:**
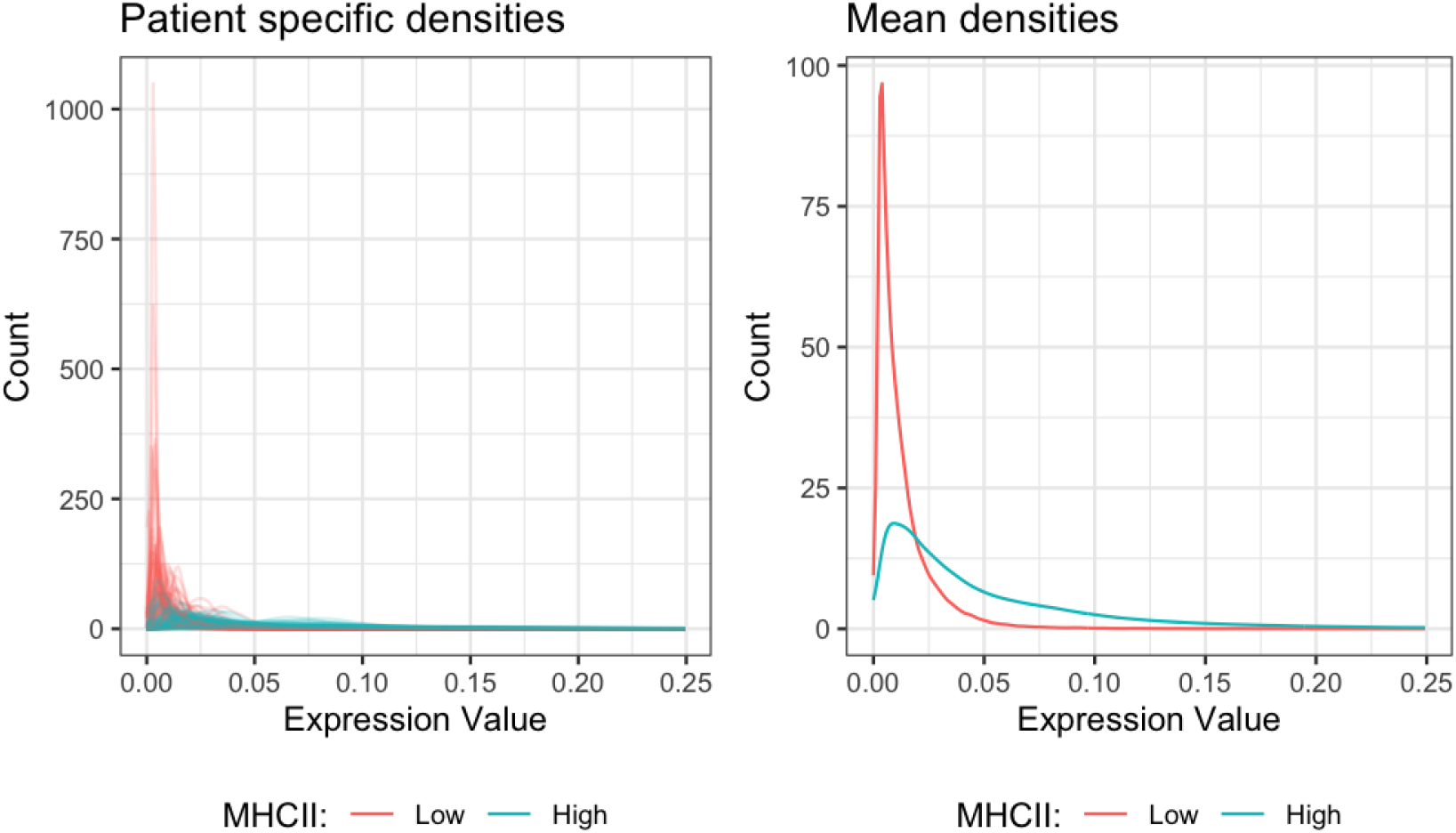
Individual (on the left) and mean (on the right) HLA-DR marker probability density (in CK+ tumor cells) of the subjects from two groups, MHCII: High and MHCII: Low. Notice that some of the subjects from MHCII: High group have density functions similar to the average of MHCII: Low group. It means that the grouping is not fully capturing the density differences between the subjects.

We also used the test based on Cox PH model with random effects from Section 2.2.3 in this case. The estimated variance of the random effect was 0.38. Following Therneau *and others* (2015)’s interpretation of the variance parameter in this context, we concluded that there are multiple subjects in the study with quite large relative risks, 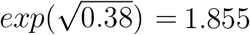 fold greater than the average subjects. However, the LRT based on integrated partial likelihoods was not significant.

### 3.2 Application to TNBC MIBI data

The triple-negative breast cancer (TNBC) MIBI dataset has images from 41 subjects. Keren *and others* (2018) categorized these subjects into three groups: “cold”, “compartmentalized” and “mixed” based on the level of immune infiltration in the TME. We were interested in studying the density of the immunoregulatory protein markers, PD1, PD-L1, and Lag3 which have been shown to have immunological relevance (Keren *and others*, 2018; Patwa *and others*, 2021). PD1 and Lag3 are primarily expressed in immune cells and “cold” subjects have very few immune cells expressing them. Thus, we focused our analysis on 33 non-”cold” subjects. For PD1 and Lag3, we studied their density only in immune cells of a subject and for PD-L1 we studied its density both in immune and tumor cells of a subject. For every marker, we computed the JSD matrix between the subjects and performed a hierarchical clustering to classify the subjects into two groups as discussed in 2.2.2. Then, we tested the vector of cluster labels for association with two available outcomes: recurrence and survival. Figure 4 shows the Kaplan-Meier curves corresponding to the three markers for both survival (left column) and recurrence (right column). We noticed that the HR of survival was large (HR = 2.824) and significant (*p* < 0.0346) for PD1 marker, indicating that the differences in PD1 marker density is associated with risk of death. For PD1, the HR of recurrence was large as well (HR = 2.065) but was not significant. For other two markers, we did not find any significant results (at level 0.05). However, it is worth pointing out that the HR of both survival and recurrence for PD-L1 were quite large (3.49 and 2.84 respectively), alluding to a possible association of PD-L1 marker density with both risk of death and risk of recurrence. We should also keep in mind that the sample size for this particular analysis was quite low which could limit our power.

**Figure 4:**
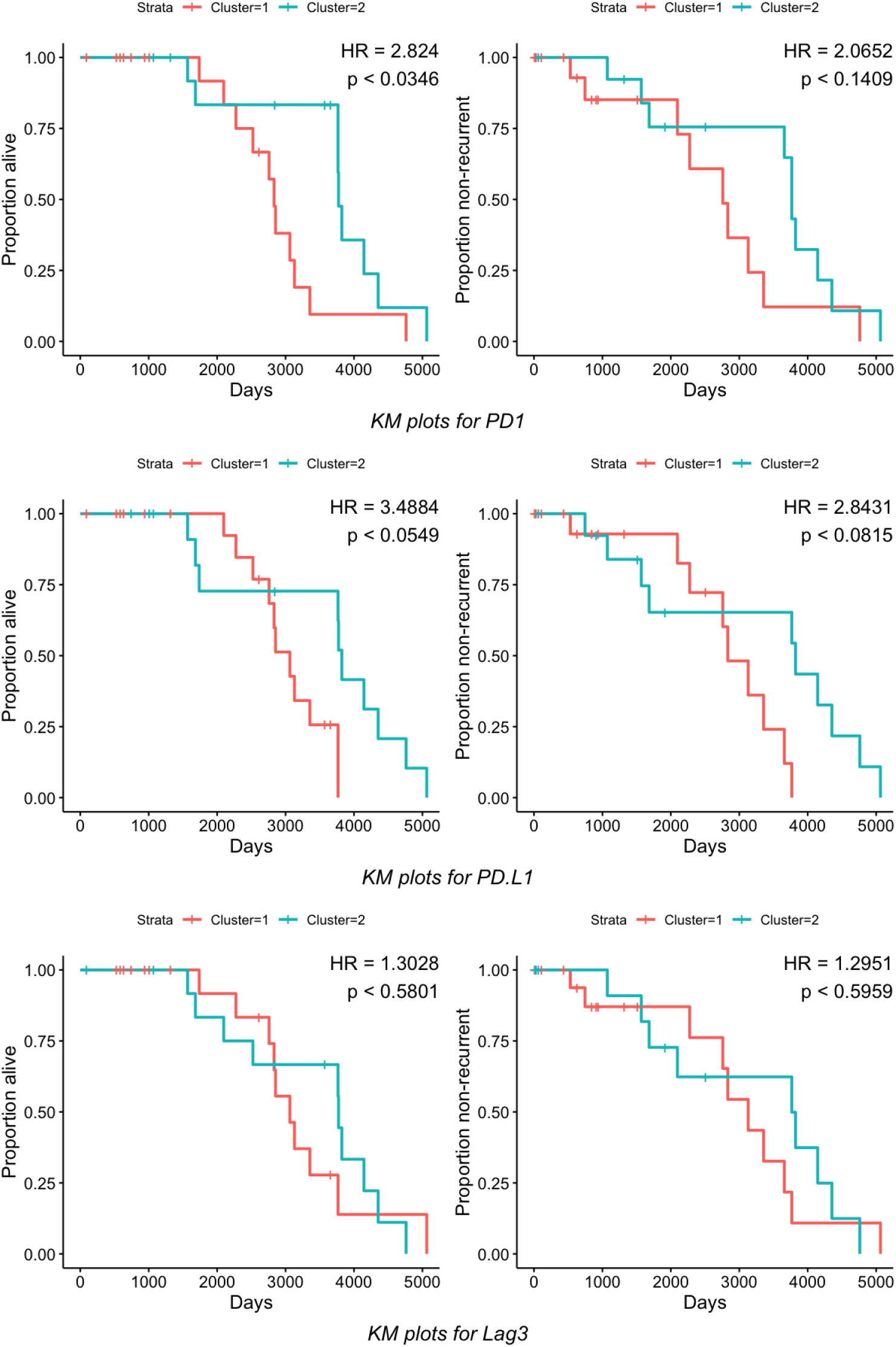
KM Plots of overall survival (left) and recurrence (right) of 33 subjects color coded by the clusters found using markers: PD1, PD-L1 and Lag3, using our method. We notice that difference in PD1 density has significant effect on overall survival.

### 3.3 Simulation study application

Next, we assessed the performance of JSD based clustering (from Section 2.2) in different simulation setups. We tried to replicate the characteristics of the real dataset discussed in Section 3.1. In Figure 2, we showcased the mean of HLA-DR probability densities of the subjects from the two clusters identified by JSD based clustering method. We found that these mean densities can be well approximated using Beta distributions (Gupta and Nadarajah, 2004) with different set of parameters (*α*, *β*). To find out the set of parameters (*α*, *β*) that would approximately replicate the mean densities of the two clusters observed, we considered the following strategy. To replicate the mean density of cluster 1, we first computed its empirical mode, say *m*_1_. We wanted to find parameters *α*_1_, *β*_1_ so that Beta(*α*_1_, *β*_1_) had the same mode and a density function very similar to the empirical one. Matching the modes implies, 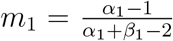. For a given value of *β*_1_, *α*_1_ is fixed and can be computed using the last equation. We considered multiple values of *β*_1_ and chose the one for which the simulated density appeared to be closest to the real one. We repeated the above steps for replicating the mean density of cluster 2 as well.

The modes of the mean densities of cluster 1 and 2 were respectively, *m*_1_ = 0.0039 and *m*_2_ = 0.0176. The mean density of cluster 1 was well approximated by a Beta distribution with *α*_1_ = 2.17, *β*_1_ = 300 and the mean density of cluster 2 by a Beta distribution with *α*_2_ = 1.78, *β*_2_ = 45. Refer to Figure 5 and 6 to check how well the real and simulated densities agree. Finding the suitable sets of parameters of Beta distribution that best summarized the real data mean densities, we focused on two different simulation studies next.

**Figure 5:**
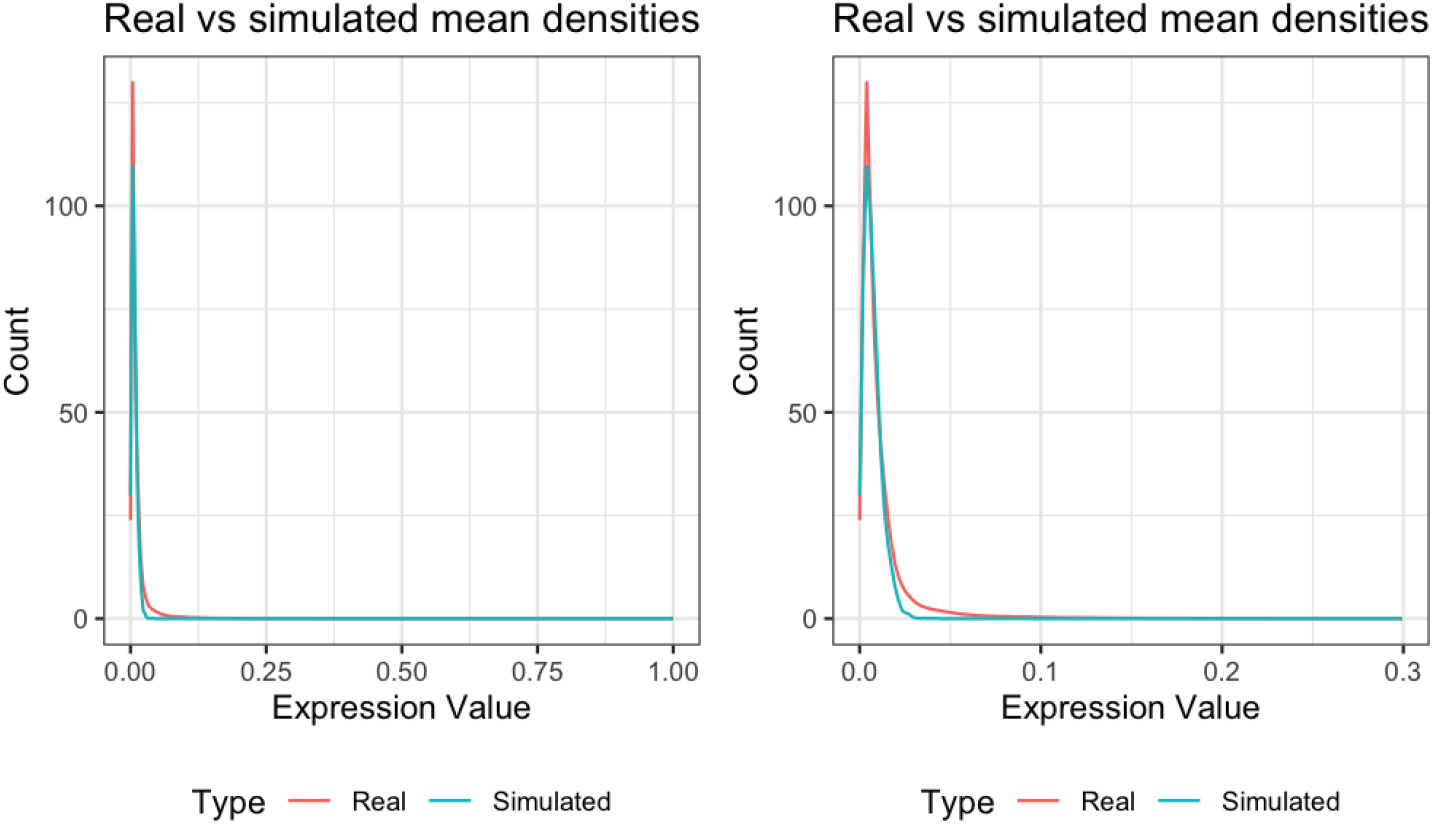
Comparing the probability density of Beta(2.17, 300) to the real mean density of cluster 1. On the left, are shown the densities on the entire range of expression value: (0, 1). On the right, we zoom into the lower expression values and the same densities are shown only between (0, 0.3). Even though there appears to be a difference in the modes of the densities, their overall shapes are quite close.

**Figure 6:**
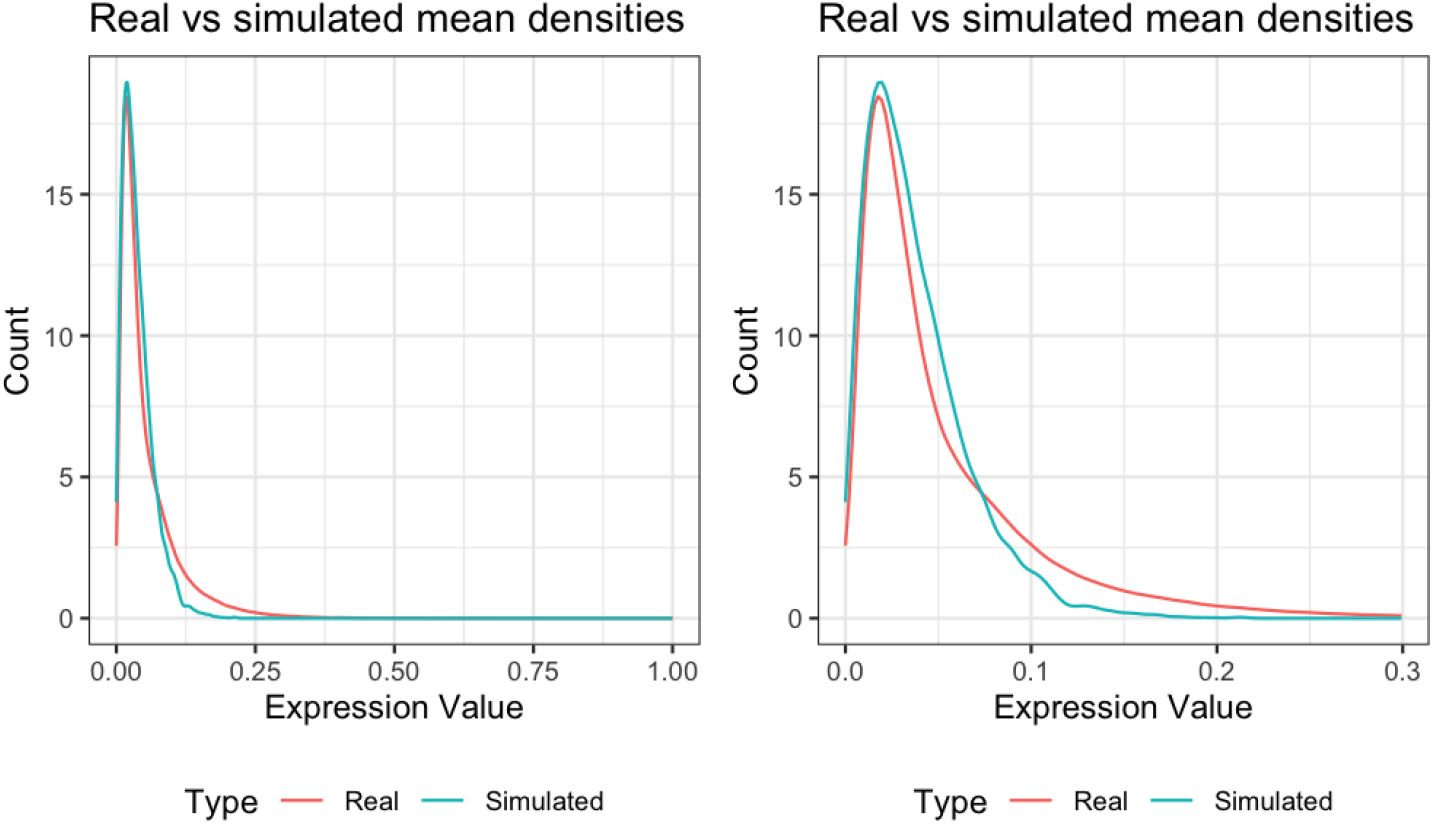
Comparing the probability density of Beta(1.78,45) to the real mean density of cluster 2. On the left, are shown the densities on the entire range of expression value: (0, 1). On the right, we zoom into the lower expression values and the same densities are shown only between (0, 0.3). The overall shapes of the densities are quite similar.

#### 3.3.1 Simulation with densities close to the real mean density of cluster 1

We assumed that there were two groups with *N*_1_ and *N*_2_ subjects (*N* = *N*_1_ + *N*_2_). We considered *N*_1_ = 60, *N*_2_ = 40. We assumed that each subject *j* had same number of cells i.e. *n_j_* = *n*. Two values of n: 200 and 2000 were considered. The marker data for a cell of a subject from group 1 was simulated from Beta(2.17, 300) i.e. the distribution which best summarized the real mean density of cluster 1. The marker data of a subject from group 2 was simulated from Beta(*x*, 300) where x is such that the mode of this distribution was higher than 0.0039 by a percentage of *l* i.e. *x* satisfied

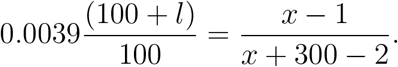

Five different values of *l*: 10, 20,100,150 and 200 were considered. We wanted to study how well JSD based clustering approach can classify the subjects into their respective groups. We used two measures: adjusted Rand index (ARI) (Santos and Embrechts, 2009), adjusted mutual information (AMI) (Romano *and others*, 2014) which are popular in semi-supervised learning literature. We compared our method with the thresholding based approach described in Section 2.1. As discussed earlier, the thresholding based approach requires two thresholds *t*_1_ and *t*_2_. Since, we did not know what thresholds would possibly be suitable in this simulation setup, we varied *t*_1_ between 95% and 97.5% quantiles of the full marker data (concatenating marker data of all the subjects) and kept *t*_2_ at 0.01. These two methods were referred to as 95% and 97.5% thresholding respectively. Table 2 lists the performance of all these methods. We noticed that when the number of cells and difference in modes were both small (*n* = 200, *l* = 10), all the methods performed poorly in terms of ARI and AMI. However, the performance of JSD based clustering improved hugely when the number of cells increased (*n* = 2000). Even for a moderate difference in modes (*l* = 50), JSD based clustering achieved close to 1 accuracy, whereas thresholding methods kept achieving little to zero accuracy.

**Table 2:** Performance of different methods in the simulation with densities close to the mean density of cluster 1 as described in Section 3.3.1. JSD based clustering performs systematically better than the thresholding approaches in all the cases. When the number of cells is large, JSD based clustering performs well even for small differences in modes.

| Measure of performance | Number of cells | Percentage difference in modes | JSD based clustering | 95% thresholding | 97.5% thresholding |
| --- | --- | --- | --- | --- | --- |
| ARI | $n = 200$ | 10 | 0.0744 | 0.0014 | 0.0234 |
|  |  | 20 | 0.3988 | 0.0029 | 0.0551 |
|  |  | 50 | 0.9808 | 0.0236 | 0.2264 |
|  |  | 100 | 1.0000 | 0.1979 | 0.6324 |
|  |  | 200 | 1.0000 | 0.8628 | 0.9570 |
| | $n = 2000$ | 10 | 0.8029 | 0.0000 | 0.0000 |
|  |  | 20 | 0.9530 | 0.0000 | 0.0001 |
|  |  | 50 | 1.0000 | 0.0000 | 0.0299 |
|  |  | 100 | 1.0000 | 0.0040 | 0.8105 |
|  |  | 200 | 1.0000 | 0.9907 | 1.0000 |
| AMI | $n = 200$ | 10 | 0.0713 | 0.0026 | 0.0144 |
|  |  | 20 | 0.3344 | 0.0041 | 0.0358 |
|  |  | 50 | 0.9662 | 0.0283 | 0.1802 |
|  |  | 100 | 1.0000 | 0.2001 | 0.5421 |
|  |  | 200 | 1.0000 | 0.8088 | 0.9239 |
| | $n = 2000$ | 10 | 0.7233 | 0.0000 | 0.0000 |
|  |  | 20 | 0.9976 | 0.0000 | 0.0001 |
|  |  | 50 | 1.0000 | 0.0000 | 0.0322 |
|  |  | 100 | 1.0000 | 0.0039 | 0.7609 |
|  |  | 200 | 1.0000 | 0.9865 | 1.0000 |

#### 3.3.2 Simulation with densities close to the real mean density of cluster 2

We again considered two groups respectively with *N*_1_ and *N*_2_ subjects each of whom had *n* cells. This time, the marker data for a cell of a subject from group 1 was simulated from Beta(1.78, 45) i.e. the distribution which best summarized the real mean density of cluster 2. The marker data for a cell of a subject from group 2 was simulated from Beta(*x*, 45) where *x* is such that the mode of this distribution was higher than 0.0176 by a percentage of *l* i.e. *x* satisfied

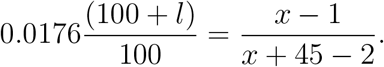

We again considered *N*_1_ = 60, *N*_2_ = 40 (and thus, *N* = 100). Two values of *n*: 200 and 2000 and five values of *l*: 10, 20, 100, 150, 200 were considered. Table 3 lists the performance of all the methods. Once again, JSD based clusetring outperformed the thresholding based approaches in all the cases. One interesting observation is that the thresholding based approaches seemed to be performing worse in this simulation setup compared to the previous one. Possibly, a different set of (*t*_1_, *t*_2_) would have been more appropriate in this scenario. It reiterates the point that the subjectivity of the thresholding based approaches can hugely alter or affect the performance.

**Table 3:**
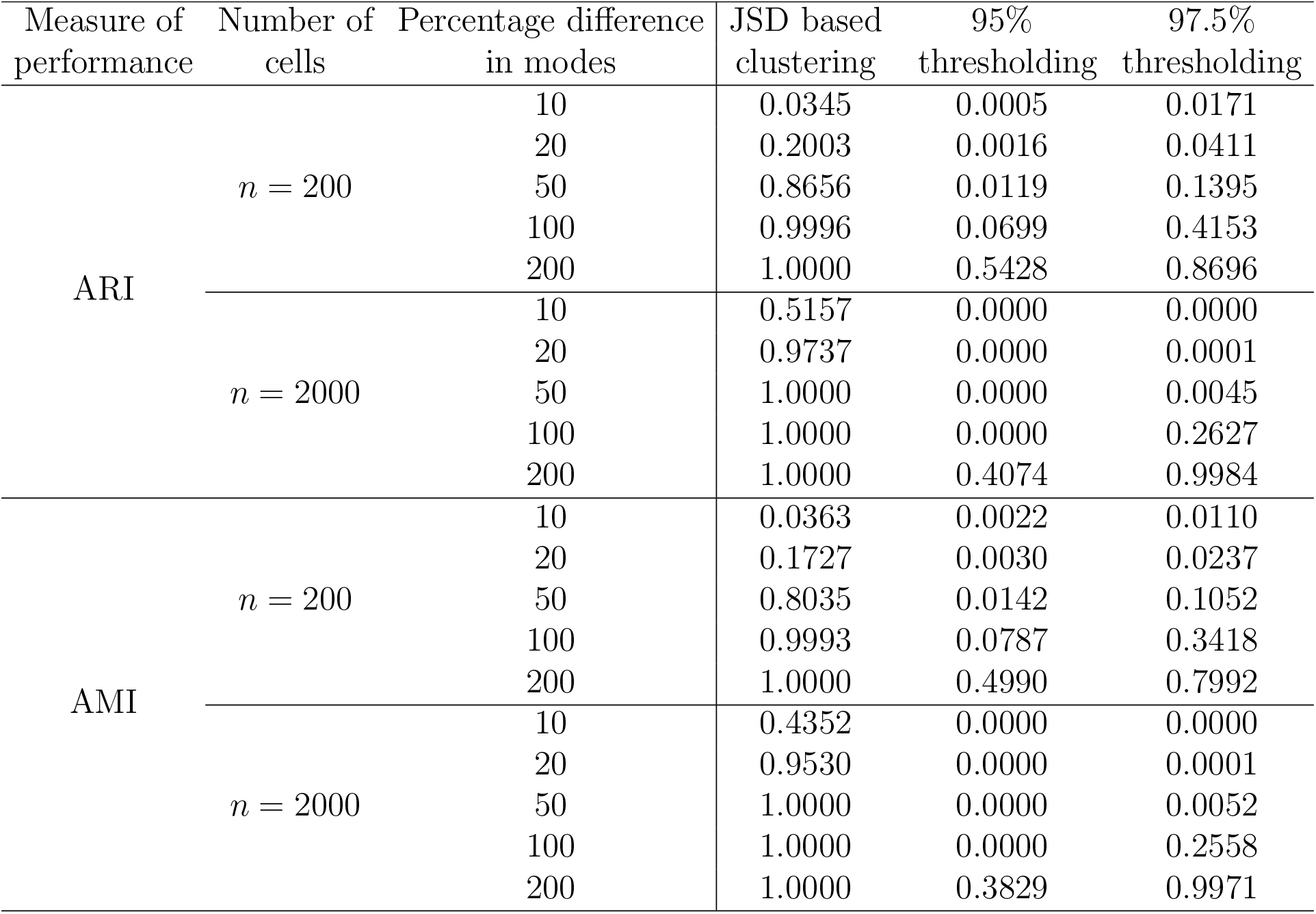
Performance of different methods in the simulation with densities close to the mean density of cluster 2 as described in Section 3.3.2. JSD based clustering performs systematically better than the thresholding approaches in all the cases. When the number of cells is large, JSD based clustering performs well even for small differences in modes.

#### 3.3.3 Simulation favoring the thresholding based approach

Next, we devised a simulation where the true values of the thresholds: (*t*_1_, *t*_2_) were known. And the marker data generation process was dependent on those. Recall that *t*_1_ controls how we define a cell to be positive for a marker and *t*_2_ controls how we cluster the subjects into two groups. The simulation strategy was as follows. We considered two groups with respectively *N*_1_ and *N*_2_ subjects, each with *n* cells. We kept *N*_1_ = 40 and *N*_2_ = 60 and varied *n* between 200 and 2000. We wanted the subjects in group 1 to have *t*_2_% positive cells and the subjects in group 2 to have more than *t*_2_% positive cells. We describe the process of simulating the marker data of the non-positive cells first. For subjects in group 1, we made sure that they had (100 – *t*_2_)% non-positive cells by randomly choosing (100 – *t*_2_)*n*/100 cells out of the total of *n*. Let 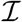 denote the set of indices of those non-positive cells for subject *j*. Next, the marker data of *i*-th cell from set 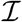, *X_ij_* was simulated from Beta(2.17, 300). To avoid any notational confusion, we highlight that *X_ij_* can be thought of as *X_kij_* from the methods section. Since we were dealing with a single marker, we dropped the index *k* for simplicity. Once, all the 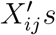 were generated, the values were scaled to be in the range (0, *t*_1_) using the transformation, 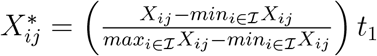 for 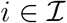. Next, we describe the process of simulating the marker data of the positive cells. The marker data of the positive cells (i.e. *X_ij_*’s for 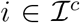) were again simulated from Beta(2.17, 300) and a constant of *t*_1_ was added to them, 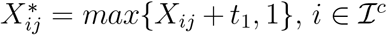. Thus, we had generated whole cell-level data of a subject *j* from group 1, 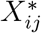 for *i* ∈ {1,…, *n*} making sure there were only *t*_2_% cells having marker expression more than *t*_1_.

For subjects in group 2, we had to make sure that they have more than *t*_2_% positive cells. So, for such a subject *j* we simulated a number, 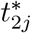 from Uniform(*t*_2_, 0.9) and repeated all the steps used in simulating group 1 with 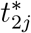 in place of *t*_2_. Note that for both the groups, we used Beta(2.17, 300) to simulate the initial cell-level data (*X_ij_*) and then slightly transformed it 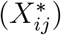 to maintain the threshold criteria. One might as well vary the primary distribution as well between the groups but our goal was to create the hardest possible simulation scenario for our method where there would be no explicit difference in marker ensities between two groups. We considered two different values of *t*_1_: 0.05, 0.1 and five different values of *t*_2_: 0.005, 0.01, 0.05, 0.1 and 0.2. Table 4 lists the performance of JSD based clustering for all combinations of the parameters. We found out that our method performed better for higher values of *t*_2_. The value of *t*_1_ and the value of n did not have any apparent impact on the performance. Keep in mind that using the thresholding approach in this simulation setup with the known values of (*t*_1_, *t*_2_) one would achieve ARI and AMI accuracy of 1 in all the cases. However, as we have repeatedly pointed out, knowing the true values of (*t*_1_, *t*_2_) will never be possible in real data.

**Table 4:** Performance of JSD based clustering in the simulation from Section 3.3.3. The method performs better for larger values of *t*_2_, whereas *t*_1_ does not seem to affect the performance.

| Number of<br>cells | Measure of<br>performance | $t_2 :$ | 0.005 | 0.01 | 0.05 | 0.1 | 0.2 |
| --- | --- | --- | --- | --- | --- | --- | --- |
| $n = 200$ | ARI | $t_1 = 0.05$ | 0.760 | 0.791 | 0.801 | 0.938 | 1.000 |
| | | $t_1 = 0.1$ | 0.784 | 0.727 | 0.815 | 0.957 | 0.987 |
| | AMI | $t_1 = 0.05$ | 0.764 | 0.772 | 0.756 | 0.922 | 1.000 |
| | | $t_1 = 0.1$ | 0.712 | 0.719 | 0.765 | 0.943 | 0.987 |
| $n = 2000$ | ARI | $t_1 = 0.05$ | 0.778 | 0.727 | 0.800 | 0.936 | 1.000 |
| | | $t_1 = 0.1$ | 0.784 | 0.727 | 0.808 | 0.965 | 1.000 |
| | AMI | $t_1 = 0.05$ | 0.733 | 0.678 | 0.755 | 0.919 | 1.000 |
| | | $t_1 = 0.1$ | 0.734 | 0.680 | 0.762 | 0.951 | 1.000 |

## 4 Discussion

In multiplexed tissue imaging datasets, it is often of interest to stratify the subjects based on the profile of functional markers for the purpose of risk assessment (e.g. risk of recurrence, risk of death etc.). The most common approach of grouping the subjects into meaningful clusters is a thresholding-based method which requires elaborate tuning of two or more thresholds. In consequence, the method remains largely subjective and varies from one researcher to another based on their interpretation of the data. In this paper, we have developed a threshold-free method for classifying subjects based on the probability density of the functional markers in the tumor microenvironment (TME). The method is easy to interpret and free from the subjectivity bias.

In our method, we treat the expression of a functional marker in a subject as a continuous random variable and compute its kernel density estimate based on its observed value in the cells of the TME. Once the marker density estimates for all the subjects have been computed, we use the Jensen-Shannon distance to quantify the difference in marker densities between the subjects. If the distance between two subjects is large, it means that they have very different marker expression profiles. Next, the computed distance matrix is used in either of the following two ways. It can be subjected to hierarchical clustering to group the subjects into clusters and the cluster-labels can be tested for association with outcomes of interest (e.g. recurrence, survival). Or it can be used directly in a linear mixed model setup for testing association with outcomes of interest.

We analyzed two highly complex multiplex tissue imaging datasets, an mIHC lung cancer dataset from University of Colorado School of Medicine and a publicly available triple negative breast cancer MIBI data. In the lung cancer dataset, we found out that the difference in HLA-DR marker density between subjects was significantly associated with their 5-year overall survival. In the breast cancer dataset, we found out that the difference in the density of immunoregulatory protein PD-1 was associated with the overall survival. Next, we replicated the characteristics of the lung cancer dataset in two simulation scenarios and showcased the robustness of our method in comparison with the thresholding-based method. In the final simulation setup, we aimed to simulate a dataset favoring the principles of the thresholding method. We showed that our method performed competently even in that scenario.

In this paper, we have focused on analyzing each of the functional markers separately. Our next goal will be to study the joint effect of multiple functional markers. One naive way of studying the joint effect would be to sum up the distance matrices corresponding to different functional markers creating a new distance matrix. This aggregated distance matrix would capture the overall difference in densities of the different markers. However, the approach is essentially assuming that the markers are independent and will be incapable of capturing complex interplay between the markers. In that light, one possible alternative would be to compare multivariate probability density of the markers across different subjects which, on the other hand, can turn out to be extremely computationally demanding. Therefore, we would study all these approaches in much greater details as a part of our next work. Additionally, we would further validate the applicability of our method using datasets coming from other imaging platforms, such as CODEX (Goltsev *and others*, 2018) and Visium (Tippani *and others*, 2021).

## 5 Software

Software in the form of a GitHub *R* package, together with an example data-set and complete documentation is available at this link, https://github.com/sealx017/DenVar.

## Acknowledgments

We thank the Human Immune Monitoring Shared Resource and support of the University of Colorado Human Immunology and Immunotherapy Initiative for their expert assistance in multiplex IHC and generation of the ovarian and lung datasets. We acknowledge the support of the University of Colorado Cancer Center Support Grant (P30CA046934). S.S. is funded by the Grohne-Stepp Endowment from the University of Colorado Cancer Center.

## Notes

### Competing Interest Statement

The authors have declared no competing interest.

